# Separating feedforward and feedback dynamics using time-frequency-resolved connectivity: a hybrid model of left ventral occipitotemporal cortex in word reading

**DOI:** 10.1101/2025.06.20.660651

**Authors:** Jiaxin You, Olaf Hauk, Riitta Salmelin, Marijn van Vliet

## Abstract

Left ventral occipitotemporal cortex (vOT) is crucial in reading, yet its functional role is viewed either as a prelexical feedforward hub or a bidirectional interface between sensory and higher-order linguistic systems. The two competing views have usually been considered mutually exclusive. To comprehensively explain the functional complexity of left vOT, we investigated the temporal and spectral dynamics of information flows involving left vOT during visual word and pseudoword reading using magnetoencephalography (MEG) and two directed connectivity metrics, i.e., phase slope index (PSI) and Granger causality (GC). Specifically, feedforward connectivity from low-level visual areas to vOT was observed for all conditions, with the strength of orthographic information flow to left superior temporal cortex (ST) modulated by stimulus word-likeness. Conversely, feedback flow from left ST to vOT appeared for pseudowords that allow top-down linguistic constraints to facilitate reading, and occurred later for word-like than complete pseudowords. Our findings suggest that left vOT during word reading functions in a hybrid manner: operating in efficient feedforward mode for familiar word recognition while flexibly recruiting bidirectional processing for unfamiliar pseudowords. By disentangling feedforward and feedback dynamics with high temporal and spectral resolution, we empirically reconcile competing theories of vOT function.

## 1 Introduction

Word reading relies critically on left ventral occipitotemporal cortex (vOT) [1, 2], but its precise functional role remains contentious [3]. One perspective considers the left vOT to be involved in prelexical orthographic processing within a hierarchical model of visual word reading [4]. An alternative view proposes that it integrates bottom-up sensory inputs and top-down feedback from higher-order linguistic regions [5].

Numerous studies have investigated the bottom-up and top-down influences on left vOT during visual word recognition. Using fMRI, primarily feedforward information flows involving left vOT were identified from lower-level visual cortex to left vOT [6] and from vOT to higher-order linguistic regions [7]. Furthermore, Li et al. [8] suggested that the increased functional connectivity between left vOT and superior temporal gyrus might reflect feedback modulation, but they provided no direct evidence of its directionality. Notably, the low temporal resolution of fMRI may constrain its ability to clearly distinguish between bottom-up and top-down information flows, which requires information about temporal relationships [9].

Electro- and magnetoencephalography (EEG and MEG) are powerful tools for investigating the fast temporal dynamics of functional brain interactions [10, 11]. For example, using MEG, [12] reported a stronger feedback connection from left inferior frontal cortex to vOT for words than false fonts in the first 200 ms. Intracranial EEG studies also showed early top-down feedback before 150 ms [13], and later phase-locking with more anterior areas from 170 to 400 ms [14]. Moreover, Whaley et al. [15] found bidirectional influences linking the left frontal and sensorimotor region with left vOT between 50 and 600 ms. These findings collectively underscore the dynamic temporal information flow involving left vOT. Still, the evidence has not always been consistent. To provide more fine-grained insights, the present study sought to examine whether and how information flows in left vOT are modulated by different levels of word-likeness.

Rhythmic synchronization has been proposed as a possible mechanism of facilitating inter-regional information flow [16]. In the visual system, signals at gamma band frequencies were found to subserve feedforward and alpha/beta rhythm feedback influences [17, 18]. However, during language processing, feedback connectivity was observed at beta band frequencies [19, 20, 21] and feedforward connectivity at alpha band frequencies [20] or high gamma band frequencies [19]. We were interested in whether such frequency-specific directed connectivity mechanisms also hold for left vOT involved in visual word reading. Alternatively, it is possible that the single-word recognition process may not be exclusively captured by oscillatory activity in canonical frequency bands. Therefore, we need analysis methods with both high temporal and spectral resolution to track those processes.

In the present study, we exploited our earlier MEG dataset recorded during reading of words and pseudowords with different levels of word-likeness [22], created by replacing up to 3 letters of a real word. This dataset is potentially well suited to address the role of vOT in merging bottom-up and top-down influences in reading by comparing real words and unfamiliar letter strings [6]. We employed two directed connectivity measures, phase slope index [PSI; 23] and Granger causality [GC; 24], based on time-frequency representation using wavelet analysis. PSI was used to capture the overall picture of dominant unidirectional information flows across the whole cortex. In contrast, GC provided a more fine-grained analysis of pairwise bidirectional interactions between left vOT and lower-order visual and higher-order language regions. By linking functional connectivity dynamics to the different levels of pseudowords, this study sought to disentangle whether word processing in left vOT was sequential (feedforward-only), bidirectional (feedforward and feedback), or hybrid–varied with the word-likeness of pseudowords, and whether the directed information flow was frequency-specific.

## 2 Methods

### 2.1 Preprocessing

Spatiotemporal signal space separation (tSSS) was applied to suppress signal interference from outside of the head [25]. The data was band-pass filtered at 0.1–40 Hz. Artifacts associated with eye movements, eye blinks and heartbeats were removed using independent component analysis (ICA). Artifact-related components initially automatically detected and then manually selected were excluded from the data. We extracted epochs from-200 to 1100 ms with respect to each stimulus presentation, including a -200–0 ms pre-stimulus baseline. Epochs with excessive peak-to-peak signal amplitudes were removed, with the cutoff threshold of 3000 fT/cm for the gradiometer sensors and 4000 fT for the magnetometer sensors.

### 2.2 Source analysis of MEG evoked responses

Field spread significantly constrains the utility of connectivity measures derived from sensor-level data [26]. Therefore, we focused our investigation on neural communication at the source level. We used a surface-based cortical source space with 2562 vertices per hemisphere. We calculated the inverse operator for each participant with a loose orientation constraint of 0.2 and a depth weighting parameter of 0.8. We then performed source reconstruction based on single-trial epochs of each condition using noise-normalized dynamic Statistical Parametric Maps [dSPMs; 27]. We then applied each participant’s inverse operator to single epochs for each condition with a regularization parameter *λ*^2^ = 1, corresponding to an assumed signal-to-noise ratio (SNR) of 1. This is the recommended value in the MNE-Python framework for source estimation on epoched data, as it prioritizes noise suppression during trial-wise source reconstruction, thereby enhancing the robustness of subsequent connectivity estimation. Only activity estimated to be oriented perpendicular to the cortical surface was retained. The source estimates for each participant were morphed to a standard template brain, “fsaverage”. We applied a cortical parcellation including 139 parcels based on the Destrieux Atlas [28]. Time courses were extracted from source-reconstructed evoked responses of single trials for each condition and each subject, and inserted in the functional connectivity analysis.

### 2.3 Regions of interest

Left vOT was taken as the seed region of interest (ROI) in our connectivity analyses. To examine its interactions with low-level visual areas and high-level phonological/semantical areas, four parcels were chosen as target ROIs to represent these functionally specialized brain regions in the left hemisphere: primary visual (PV), superior temporal (ST), precentral (pC), and anterior temporal (AT) cortices. All ROIs are shown in Figure 1a. The left ST and pC were chosen based on our earlier analysis of evoked responses on this same data set [22]. Left ST is typically reported to subserve semantic processing [29, 30] and pC phonological processing [31, 32]. Left AT was included as ROI due to its proposed role as a general hub region for semantic processing [33]. The PV region engaged in lower-level visual processing was included to quantify connectivity due to bottom-up perceptual input to vOT.

**Figure 1:**
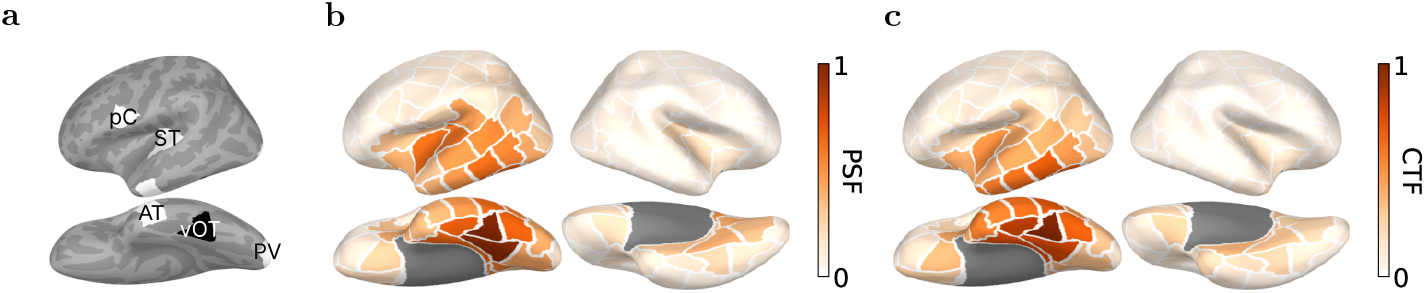
**a**, Regions of interest. The seed region is vOT and high- and low-order target regions selected for analysis pC, ST and AT. **b-c**, Source leakage patterns from left vOT, i.e., PSFs (**b**), and to left vOT, i.e., CTFs (**c**).

### 2.4 Source leakage in left vOT

Source leakage remains a concern even when the analysis is performed in the source space. To characterize the source leakage patterns regarding left vOT, we used point spread and cross-talk functions [PSFs and CTFs; 34, 35]. Following Rahimi et al. [36], we estimated the leakage indices from left vOT to all other parcels using PSFs (Figure 1b) and vice versa using CTFs (Figure 1c). These patterns were normalized against the self-leakage within left vOT. The left vOT mainly produces and receives the leakage in its vicinity, with outgoing leakage more pronounced than incoming. Specifically, left vOT predominantly leaked activity to left lingual gyrus. In contrast, incoming leakage was high only from a few adjacent parcels, which diminished progressively with distance. These findings provide a basis for evaluating the degree to which leakage may affect the estimated connectivity patterns.

### 2.5 Directed functional connectivity metrics

As MEG data are intrinsically dynamic, all connectivity measures were derived from time-frequency representation of time courses using Morlet wavelets. The analysis was performed over the entire epoch duration, spanning from -200 to 1100 ms relative to stimulus onset at 10-ms steps. To ensure adequate spectral resolution, the analysis was conducted across a frequency range of 4 to 40 Hz in 1-Hz steps. This frequency range covered canonical frequency bands from theta to low gamma. The number of wavelet cycles was set to half of each frequency value, which ensures an optimal trade-off between frequency and time resolution, which is particularly valuable for studying rapid connectivity changes during the task. We used two directed functional connectivity metrics, PSI and GC, to quantify inter-areal interactions based on the time-varying spectra obtained from continuous wavelet transform (CWT). PSI estimates whether there is a consistent positive or negative latency difference between activations in pairs of regions. Thus, it allows estimating directed connectivity values between a seed region (here vOT) and the whole brain at a relatively low computational cost. However, it does not provide information about causal relationships between time courses, and in particular it cannot disentangle bidirectional relationships [37]. In contrast, GC can deal with such causal and bidirectional relationships [38], also taking into account that every region consists of multiple partially correlated time courses (one per vertex). Thus, these two approaches complement each other well in the analysis, and allow us to assess the reliability and robustness of the connectivity results. The overall schematic is shown in Figure 2.

**Figure 2:**
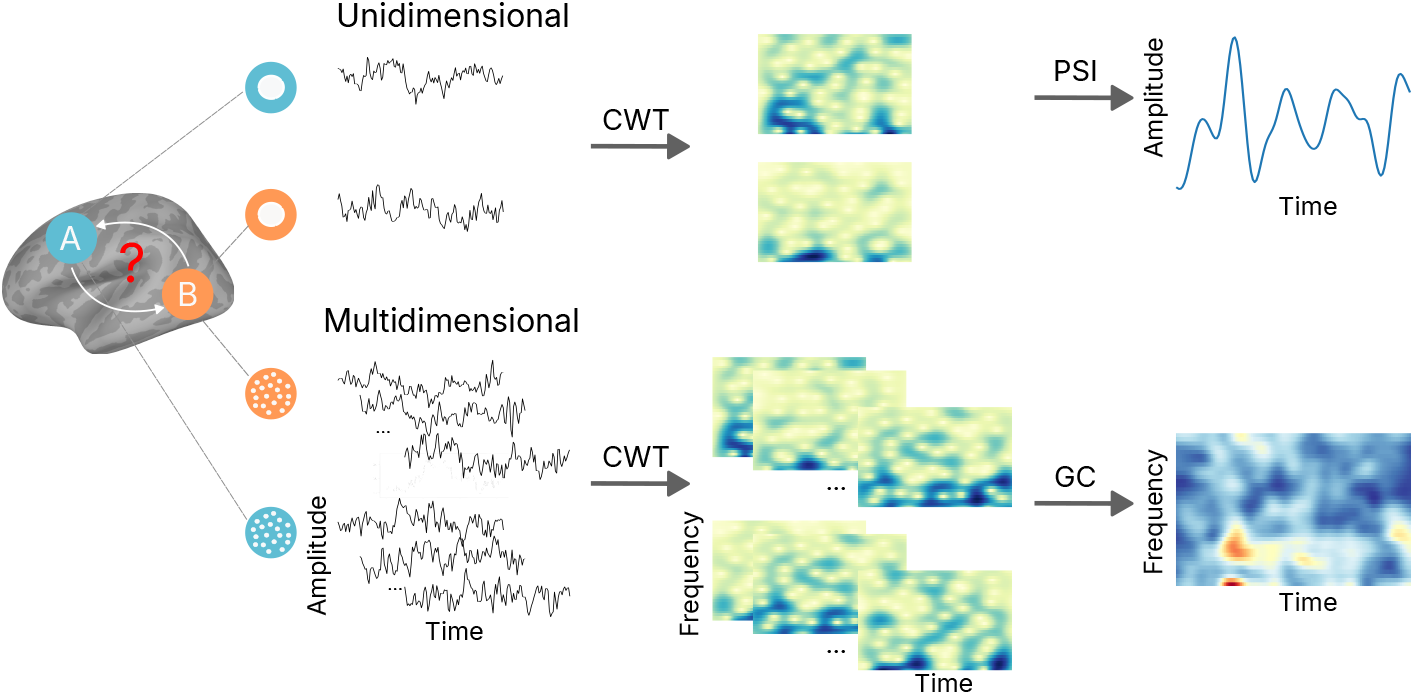
Schematic of the time-varying directed connectivity methods, phase slope index (PSI) and Granger causality (GC), used in this study. They are based on the time-frequency representation of source-level unidimensional/multidimensional time courses (− 200–1100 ms) using continuous wavelet transform (CWT; 4–40 Hz).

PSI measures asymmetrical influence indexing unidirectional interactions. It is derived from the coherency in the frequency domain:

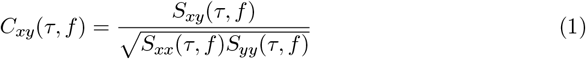

where 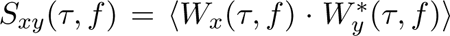 is the cross-spectrum between regions *x* and *y, W*_*x*_ is the wavelet transform of the averaged time course from region *x* and ⟨·⟩ denotes the ensemble average. *S*_*xx*_ and *S*_*yy*_ denote their respective autospectra. Using the coherency, PSI is computed as the consistency of phase differences across frequencies within a specified frequency range ℱ:

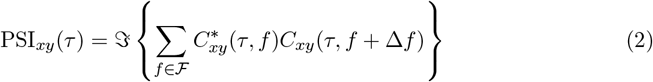

where Δ*f* is the frequency resolution, and ℑ(·) denotes taking the imaginary part. The sign of PSI indicates the lag or lead relationship between seed and target regions. We first computed the PSI between left vOT and each other parcel to provide an overall picture of the whole-cortex connectivity. To distinguish task-related connectivity from background connectivity, the measured connectivity was baselined to the mean over the baseline period (−200–0 ms) as done by Wei et al. [39]. To capture temporal evolution, we averaged the connectivity patterns within three time windows: early (100–400 ms), middle (400–700 ms), and late (700–1100 ms). We conducted separate analyses for each canonical frequency band: theta (4–7 Hz), alpha (7–13 Hz), low beta (13–20 Hz), high beta (20–30 Hz), and low gamma (30–40 Hz) [28]. However, it is not guaranteed that word recognition is reflected in narrow-band “oscillations”. Thus, this analysis was also applied at broadband 4–40 Hz. Note that frequencies below 4 Hz were not included because of our limited epoch length. We compared the results obtained with canonical frequency bands and broadband to identify important patterns that might be overlooked in narrow-band analyses and vice versa.

GC is a bidirectional connectivity metric based on state-space models. GC quantifies whether the past activity of one time series can predict the activity of another, and vice versa. Beyond measuring the dominant information flow, GC thus allowed us to track bottom-up and top-down information flows separately. Here, GC was applied on vertex-level time courses from parcel pairs to utilize multidimensional dependencies between regions. To alleviate data redundancies at the vertex level, we used principal component analysis (PCA) to reduce the data dimensions of time courses in paired regions. The projection rank for seed and target data, ranging from 5 to 7 per parcel pair, was set the same to avoid biases in connectivity estimates. Granger causal influence of *y* on *x* in the time-frequency domain is quantified by comparing the autospectrum *S*_*xx*_(*τ, f*) with its intrinsic counterpart, excluding the causal contribution of *y*, which is formulated as

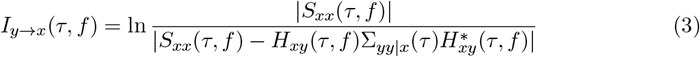

Where ***H***(*τ, f*) is the transfer matrix and Σ is the covariance matrix of the residuals of the full vector autoregressive (VAR) models. Given those, the spectral matrix is ***S***(*τ, f*) =***H***(*τ, f*)**Σ**(*τ*)***H***^*\**^(*τ, f*). We set the VAR model order as 20 for GC estimation, which yielded robust results at a moderate computational cost. In this sense, if y causally contributes to x, the numerator is greater than the denominator, i.e., Granger score *>*0. However, the original GC can yield spurious information flows due to other (uninteresting) asymmetries such as different SNRs [40]. Time-reversed GC has been proposed as a solution to this issue [41]. Practically, this was achieved by transposing the autocovariance, required for VAR models, to mimic the reversal of the original signal in time [40]. We thus estimated GC as the difference between the original GC influence (*I*) and the time-reversed one (*Ĩ*). Two directed GCs between areas *x* and *y* were defined as

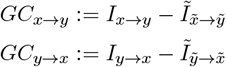

This allowed us to conservatively eliminate bias in causal influences irrelevant to cognitive processes such as noise [42]. It is worth noting that in the extreme case of unidirectional flow from *x* to *y*, the directionality of GC also flips when the signals are time-reversed 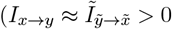 and 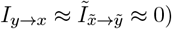 [42]. Thus, here 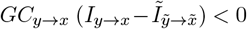 which may be counterintuitive but not surprising. Based on the asymmetry of causal interactions between hierarchical language areas, we defined the predominant or net direction of GC as

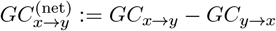

### 2.6 Statistical testing

Cluster-based permutation tests were used in this study to determine the statistical significance across subjects. For whole-cortex PSI results, we performed separate tests for three time windows (100–400, 400–700, 700–100 ms), with all parcels being entered into each test. We used 5000 permutations and a cluster-forming threshold of *t >* 1 based on a one-sample *t*-test for these tests.

For pair-wise connectivity results, the test was implemented on one-dimensional time series derived from PSI metric and on two-dimensional (time and frequency) matrices derived from GC metric. Additionally, for the GC results averaged over frequency (time-resolved) or over time (frequency-resolved), the tests were conducted on the corresponding one-dimensional series. We used 5000 permutations and a cluster-forming threshold of *t >* 2 based on a one-sample *t*-test for these tests.

## 3 Results

### 3.1 Whole-cortex directed connectivity

To obtain an overall dynamic description of bottom-up and top-down processing in left vOT, we first examined whole-cortex connectivity patterns using the PSI metric, assessing connections between each parcel and left vOT across all time windows and conditions. The grand-averaged whole-cortex PSI results from broadband analysis are shown in Figure 3a. Regions with positive PSI values (red) indicate left vOT’s leading role, while negative PSI values (blue) suggest its lagging role. Broadly, left vOT predominantly led in the early time window across all conditions but lagged later for pseudowords. During the 100– 400 ms time window, it showed strong feedforward-dominated flow (positive PSI values) to widespread cortical regions, particularly for real words and the word-like condition RL1. Conversely, significant feedback-dominated information flow (negative PSI values) appeared for less word-like RL2 and RL3 starting in the middle time windows of 400– 700 ms, while feedback for word-like RL1 occurred later, during the 700–1100 ms time window. These results suggest that the nature of information flows regarding left vOT is contingent on the word-likeness of the stimuli and changes over time. Connections with left vOT were feedforward-dominated for real words, while additional feedback-dominated flows were found for pseudowords in the late stages.

**Figure 3:**
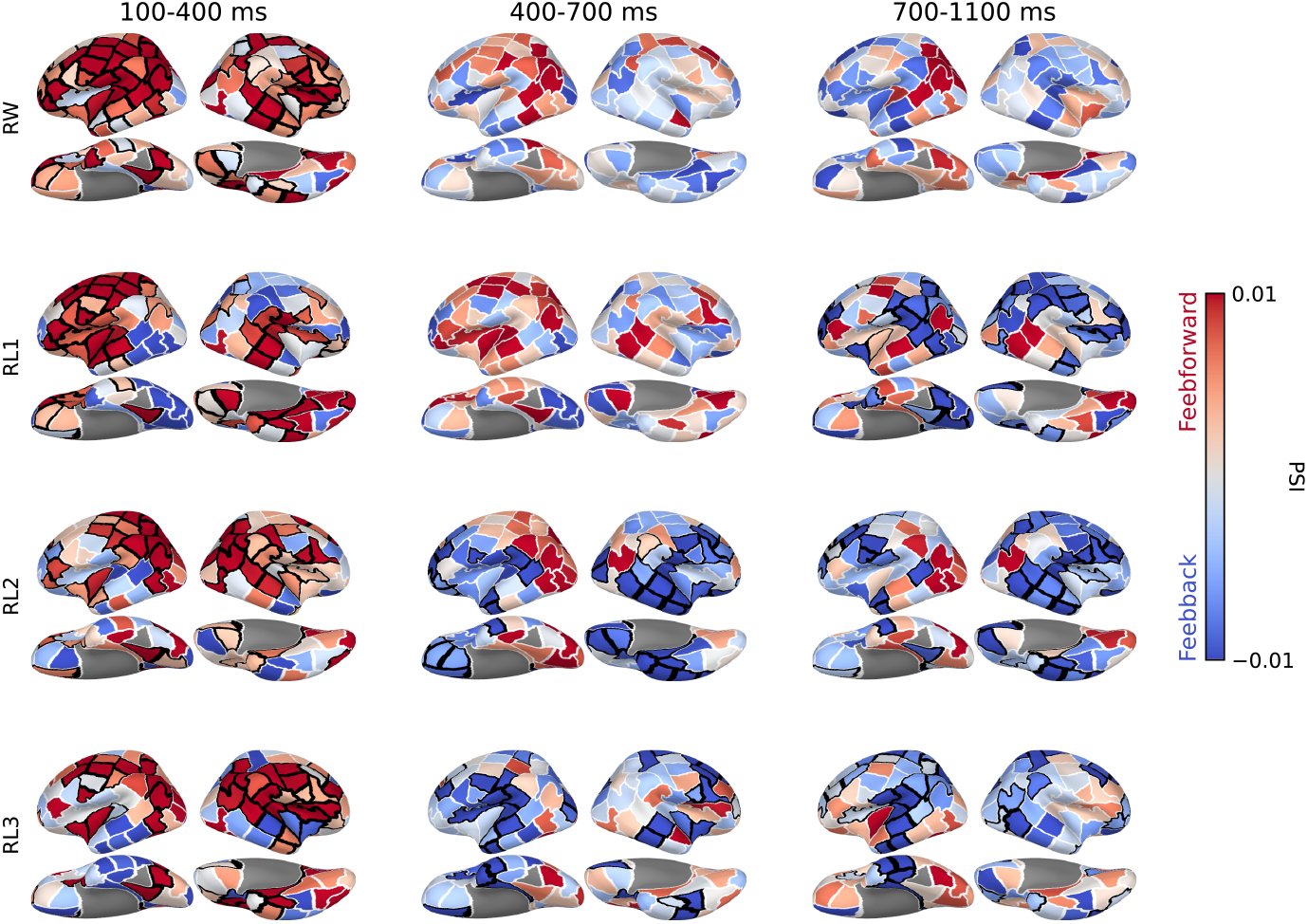
Group-level whole-cortex PSI results with left vOT as the seed region in three time windows. Each row shows the directed connectivity patterns for each condition. Regions with positive PSI values (red) indicate left vOT’s leading role, while negative PSI values (blue) suggest its lagging role. Black borders highlight the region clusters associated with *p* < 0.05 based on two-tailed cluster-based permutation tests.

To examine whether these information flows were reflected in rhythmic brain activity, we also applied the PSI analyses on separate canonical frequency bands. Results showed that the early feedforward information flow occurred in the alpha band (Figure S2), and late feedback information flow in the low beta band for RL3 (Figure S3) and in the low gamma band for RL1 and RL2 (Figure S5). However, the strengths of these flows were less pronounced compared to those observed in the broadband analysis. No significant information flow was observed in the theta or high beta bands (Figure S1, S4). These findings suggested that broadband analysis encompassed the overall picture that was captured separately in specific frequency bands. In the following, we report the PSI measures based on broadband analysis if not explicitly stated otherwise.

### 3.2 Directed connectivity with higher-order language areas

To link the bottom-up and top-down processing to the functional role of left vOT, we estimated the connectivity patterns between the left vOT and each preselected functional ROI (Figure 1a) separately. The most systematic directed connectivity measured by PSI was observed between left vOT and ST (Figure 4; see results of the connectivity with other higher-order language areas in Figure S6b and S7b). Specifically, left ST had a significant feedback influence on left vOT for RL3 (Figure 4b, orange curve), and this influence was significantly stronger for RL3 than RW(Figure 4c).

**Figure 4:**
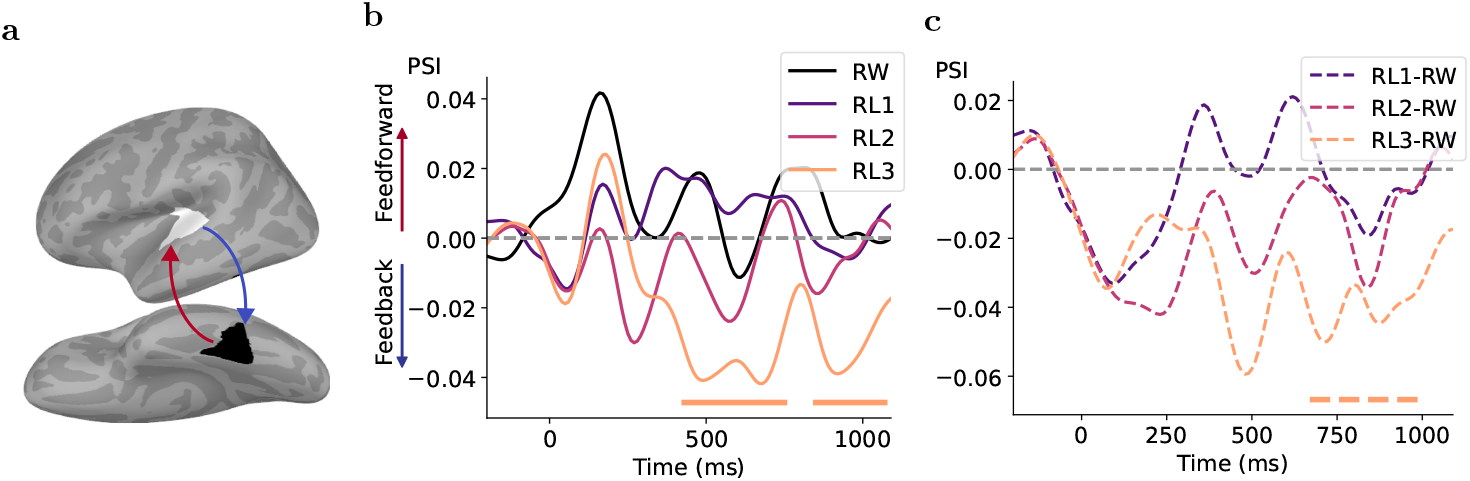
Directed connectivity patterns characterized by PSI between left vOT and ST for the different experimental conditions. **a** Schematic of information flow. Feedforward and feedback interactions are depicted in red and blue, respectively. **b–c**, Temporal dynamic of directed interactions for each condition (**b**) and their contrasts between condition pairs (**c**) from broadband analysis. Colored bars under each plot denote the time clusters associated with *p* < 0.05 based on two-tailed cluster-based permutation tests. In **b**, positive PSI values indicate the information flow from left vOT to ST (feedforward) and negative values indicate the opposite one (feedback).

For a more comprehensive characterization, we further applied GC to measure the directed connectivity patterns between left vOT and ST for each condition (Figure 5). The GC analysis reveals the time-frequency patterns of net GC, similar to PSI, but importantly also separate contributions of feedforward (vOT → ST) and feedback (ST → vOT) processes. The time-frequency maps of net GC (Figure 5a) demonstrated significant feed-forward information flow for RW and RL1, with the clusters (red) occurring around 150 ms and spanning across larger frequency ranges for RW. In contrast, another cluster (blue) was associated with feedback processing for RW around 1000 ms, spanning the theta to beta frequency bands. To track the dynamics of causal flows in the time and frequency domains, respectively, we also averaged the time-frequency GC influences separately over frequency and time to obtain time-resolved and frequency-resolved feedforward and feed-back causal influences (Figure 5a, curves on top and right in each subplot). We observed strong feedforward information flow from left vOT to ST in both time- and frequency-resolved GCs, with clusters peaking at around 150 ms and in the low-gamma frequency band for RW. However, a similar feedforward pattern was only reflected in time-resolved GC for RL1/RL2 and frequency-resolved GC for RL3. In addition, the data showed significant feedback influences, with the clusters peaking at around 900 ms for RL1 and at around 400 ms and 15 Hz for RL3. Given its intermediate word-likeness between RL1 and RL3, RL2 revealed a feedback trend between approximately 300 and 900 ms, though it did not reach statistical significance.

**Figure 5:**
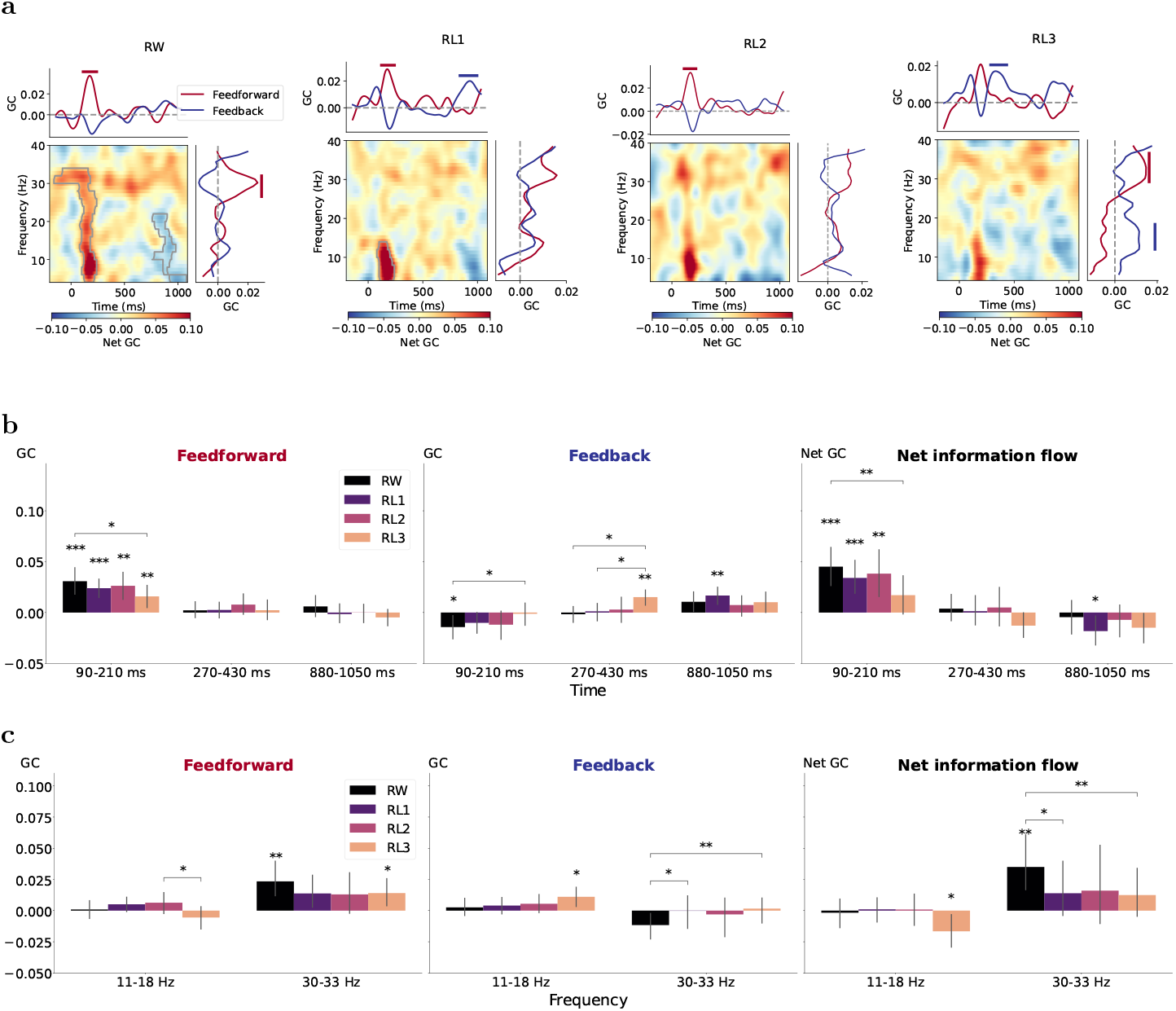
Time-varying causal influences measured by GC between left vOT and ST for each condition. **a**, In the subplot for each condition, left-bottom panel displays the time-frequency net GC map, in which significant clusters with positive values (red) indicate dominant feedforward connectivity (vOT → ST) and negative values (blue) indicate dominant feedback connectivity (ST → vOT). Curves on top and right represent separate feedforward and feedback causal influence averaged over frequency and time, respectively. Gray borders within the maps and solid bars above the plots indicate time-frequency/time/frequency clusters associated with *p* < 0.05 based on cluster-based permutation tests against zero. **b–c**, Estimated time-resolved (**b**) and frequency-resolved (**c**) GC values averaged across the selected time/frequency regions based on the clusters in **a** for feedforward direction (vOT → ST), feedback direction (ST → vOT) and net information flow from left vOT to ST. Asterisks above single bars denote the significant difference in estimated GC values from zero (one-sample t-test) and those above bar pairs denote the significant difference between conditions (paired t-test; ***, *p* < 0.001; **, *p* < 0.01; *, *p* < 0.05). In all plots, negative GC values imply a strong asymmetry of causal flow.

Since cluster-based permutation tests do not allow direct inferences about the spatial, temporal, or spectral extent of the significant clusters [43], we interpreted the clusters as time windows and frequency bands of interest for further examination. When clusters overlapped, we took their intersection. The averaged Granger causal strength for feedforward direction (vOT→ST), feedback direction (ST→vOT), and net information flow from left vOT to ST in the time windows of interest are depicted in Figure 5b and in the frequency bands of interest in Figure 5c. The feedforward processing characterized by GC was significant at 90–210 ms for all conditions and 30–33 Hz for RW and RL3. Furthermore, the feedforward information flow was stronger for RW and weaker for RL3 at 90–210 ms. In contrast, feedback flow was highlighted for RL3 at 270–430 ms and 11–18 Hz; it was significantly stronger than for RW and RL1 only in that time period. RL1 showed later feedback influence at 800–1050 ms.

Taken together, general consistency between connectivity patterns between left ST and vOT as obtained from PSI and GC was the ST→vOT feedback influence which was stronger for RL3 than RW and RL1. No consistent or systematic results were observed from the two connectivity methods between left vOT and the other preselected higher-order ROIs, i.e., left AT and pC (Figure S6 and S7).

### 3.3 Directed connectivity with lower-order visual area

To quantify how left vOT interacted with the lower-order visual area that receives the perceptual input, we measured the connectivity patterns between the left vOT and PV as shown in Figure 6. Characterized by PSI, there were peaks of feedforward-dominated information flow from left PV to vOT for all conditions at around 100 ms corresponding to the latency of visual activation, but reaching significance only for RL1 (Figure 6b, left). The directed information flows did not significantly differ between conditions (Figure 6b,right). In contrast, the time-frequency map of net GC indicated dominant feedforward interactions across all conditions, with clusters occurring around 100 ms and spanning multiple frequency bands(Figure 6c). These interactions were also reflected in the time-resolved GC curves of separate feedforward and feedback processes(Figure 6c, top in each subplot). The significant negative GC values for feedback direction suggested a strong asymmetry of causal flow (strong feedforward dominance) where the GC flipped as the data were time-reversed [42]. Figure 6d further confirmed the overwhelming feedforward sweep from left PV to vOT in the time window of 60–140 ms, which was extracted based on the clusters in Figure 6c. The rapid feedforward processing was not frequency-specific. Similar to the PSI results, the early feedforward causal flows were not modulated by stimulus conditions.

**Figure 6:**
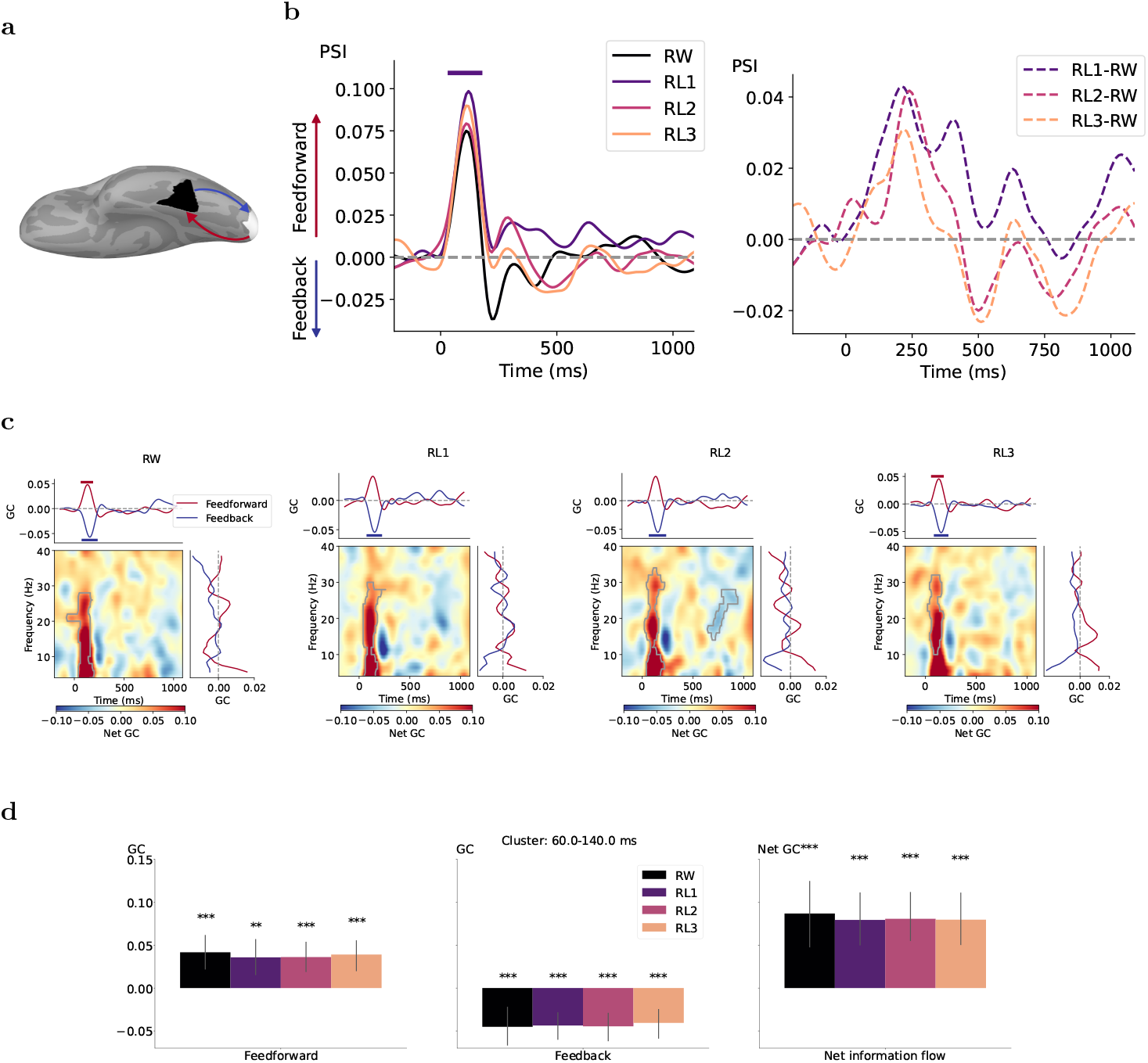
Time-varying directed interactions between left PV and vOT for each condition. **a** Schematic of information flow. Feedforward and feedback interactions are depicted in red and blue, respectively. **b**, Temporal dynamic of PSI values for each condition (left) and their contrasts between condition pairs (right). Colored bars under each plot denote the time clusters associated with *p* < 0.05 based on two-tailed cluster-based permutation tests. **c**, In the subplot for each condition, left-bottom panel displays the time-frequency net GC map, in which significant clusters with positive values (red) indicate dominant feedforward connectivity (PV → vOT) and negative values (blue) indicate dominant feedback connectivity (vOT → PV). Curves on top and right represent separate feedforward and feedback Granger causal influence averaged over frequency and time, respectively. Gray borders within the maps and solid bars above/below the plots indicate time-frequency/time/frequency clusters associated with *p* < 0.05 based on cluster-based permutation tests against zero. **d**, Estimated time-resolved GC values averaged across an early time window (extracted based on selected time clusters in **a**) for feedforward direction (PV → vOT), feedback direction (vOT → PV) and net information flow from left PV to vOT.

## 4 Discussion

The present study investigated the temporal and spectral dynamics of directed information flow to and from the left vOT to elucidate its functional role in visual word processing. By utilizing a word-pseudoword reading paradigm and employing PSI and GC connectivity measures, our findings support a hybrid model adaptively integrating early feedforward and late feedback influences on left vOT, as shown in Figure 7.

**Figure 7:**
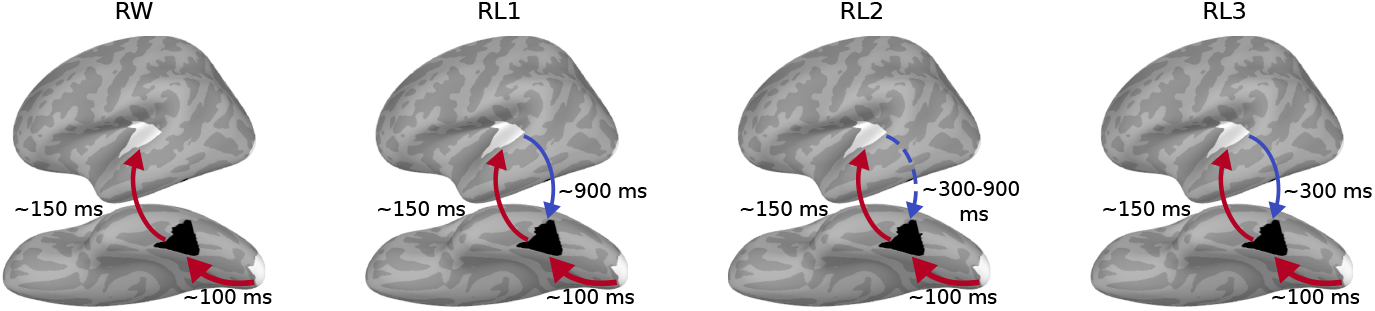
Summary schematic of directed connectivity of left vOT with higher-order ST and lower-order PV for each experimental condition. Red arrows denote feedforward flows and blue arrows denote feedback flows, along with their corresponding peak timing. Arrow thickness represents connectivity strengths, as shown in previous figures. The dashed arrow indicates a connectivity trend that did not reach statistical significance.

Directed connectivity with lower-level PV consistently exhibited strong and undifferentiated feedforward flows at around 100 ms across all conditions. This finding supports the concept of a ventral processing stream, where vOT takes in low-level visual information from PV for further (orthographic) processing. The stronger PV→vOT feedforward connectivity for pseudowords versus real words that was reported in the fMRI study [6], did not reach significance in our MEG study.

Whole-cortex PSI analysis revealed an overall picture of dominant feedforward processing for all conditions from left vOT to wide-spread anterior cortical regions in the time window of 100–400 ms. This is in line with the view that left vOT in visual word reading involves bottom-up orthographic processing [4], which occurs at around 150–200 ms in MEG studies [29, 30]. This causal flow from left vOT to ST extends the finding in a previous fMRI study [7] by incorporating temporal information. The weakest feedforward flow occurred in RL3, as demonstrated by GC analysis between vOT and ST (90–210 ms). This modulation was due to reduced bottom-up feedforward processing rather than cancellation by opposing top-down feedback. This result suggests that within vOT (or via some other input route to vOT that we are not examining here) there is early processing that distinguishes between real words and pseudowords. This supports the view that left vOT hosts a neural representation for whole real words (an orthographic lexicon) [44, 45]. The reduced feedforward flow for RL3 may be attributed to a weaker spelling-sound/semantic connection based on orthographic mapping theory [46].

After the initial feedforward sweep, feedback interactions ensued, particularly from left ST and particularly for pseudowords. Feedback influences from higher-order regions were detected largely similarly by both connectivity methods, though with some variation in the exact timing for RL1 and RL3. These results align with the feedforward view of temporal modularity, which posits that word recognition depends not only on the properties of orthographic structure but also on phonological and semantic information retrieved later [47]. Such later retrieval may enable top-down interactivity after the completion of initial orthographic processing [3]. This feedback from higher-level language areas appears necessary to resolve uncertainty in accessing the lexical entry of unambiguous pseudowords (RL1 and RL3) when strict feedforward processing alone is insufficient. Notably, RL3 showed significantly stronger feedback from left ST to vOT than the other conditions. Interestingly, RL2, representing an intermediate level of word-likeness between RL1 and RL3, only showed a feedback connectivity trend from ST to vOT. The absence of significant feedback may reflect the transitional and ambiguous nature of RL2 stimuli, which are neither complete pseudowords like RL3 nor as closely resembling real words as RL1. As a result, RL2 may fall into a representational “gray zone” where the cognitive system does not engage robust feedback mechanisms, leading to a more diffuse or inconsistent pattern that fails to reach statistical significance. This result implies that feedback strength is modulated not only by the degree of word-likeness, but also by the lexical ambiguity.

While a variety of computational models have been proposed for visual word recognition [see 48], the predictive coding framework seems particularly suited to explain these results [5, 49]; the observed feedback from vOT for pseudowords could be modeled by the prediction error stemming from the mismatch between the top-down prediction of the orthographic representation based on linguistic knowledge, which reflects the true spelling of the word, and the actual orthographic representation which contains replaced letters. RL1 pseudowords may initially be inferred into words due to their high orthographic similarity to real words, postponing the detection of mismatch and resulting in later top-down correction. Additionally, RL3, being complete pseudowords, may be rapidly identified as anomalous, prompting earlier feedback in order to resolve lexical uncertainty. These findings contrast with studies implicating feedback from higher-order regions in real-word reading [12, 13, 14]. Carreiras et al. [3] raised concerns about the theoretical necessity of top-down influences in proficient reading, and indeed our findings suggest that, in the context of single familiar word reading, left vOT primarily operates in a sequential, feed-forward mode. This is supported by the strong early feedforward connectivity from PV to vOT, and from vOT to ST, without pronounced accompanying feedback for RW.

With regard to the spectral dynamics of information flow, PSI analyses across narrow frequency bands provided no additional insights beyond broadband analyses. This result partly supports our initial speculation that rapid, low-effort, word reading may not be strongly tied to oscillatory activity within canonical frequency bands. However, whole-cortex PSI revealed some frequency-specific effects, albeit less pronounced; feedforward influences were observed in the alpha band, while feedback influences appeared in the low-beta band for RL3, mirroring a prior sentence-reading study [20]. In contrast, GC analysis revealed early bottom-up influences in the low-gamma band, consistent with prior studies on the visual system highlighting gamma rhythms’ role in feedforward processing [18, 17]. Additionally, late top-down feedback was reflected in the beta band for RL3, reinforcing the notion of beta rhythms as carriers of predictive signals. Overall, while broadband interactions dominated, frequency-specific connectivity patterns suggest that rhythmic synchronization contributes to inter-regional communication during reading [16].

Comparing the results from both PSI and GC allowed us to evaluate the robustness of our findings. Despite methodological differences, both metrics consistently identified feed-back interactions following orthographic processing for pseudoword reading, supporting a hybrid mechanism of vOT function. This effect was the most robust and reliable finding of this study. However, some discrepancies between the two methods warrant discussion. For example, GC identified significant feedforward influences from PV to vOT for all conditions as might be expected, whereas PSI only identified such influences for RL1. Similarly, GC captured feedforward sweeps from vOT to ST that PSI did not. The absence of effects from PSI is challenging to interpret and may be influenced by differences in sensitivity between the two methods. One possible explanation is that PSI, which relies on detecting consistent phase shifts across frequencies, may be more susceptible to variations in SNRs [42]. Future studies could explore strategies to mitigate these potential confounds, such as improving preprocessing techniques, increasing data length, or optimizing source reconstruction methods to enhance SNR. Additionally, PSI, as a linear method to quantify the phase difference between signals, might fail to capture nonlinear interactions [37]. While PSI can show directional coupling (i.e., the phase shift between two signals), it does not fully establish a causal relationship, which GC addresses by incorporating temporal dependencies. Additionally, GC results were more interpretable due to the use of time-reversed GC. Furthermore, connectivity patterns between other parcel pairs (vOT-AT and vOT-pC) showed no systematic results with either method. This lack of effects may be partially explained by strong mutual source leakage between vOT and AT. Though pC was far less confounded by source leakage, the lower amplitude of evoked responses in the left pC, compared to the left ST, likely weakened the connectivity strength [26]. These methodological and practical challenges underscore the need for cautious interpretation of connectivity estimates.

In summary, our results reveal distinct patterns of directed connectivity of vOT during visual word and pseudoword reading by separating feedforward and feedback influences using PSI and GC measures and exploiting the high temporal and spectral resolutions of MEG. Specifically, feedforward influences involving vOT for both word and pseudoword reading dominated early visual and orthographic processing, with the latter reflected in the low-gamma band. In contrast, late feedback from left ST emerged for pseudowords in the low-beta band and was modulated by word-likeness and lexical ambiguity. These findings reconcile competing perspectives on the left vOT function, highlighting its flexible engagement in reading under varying demands. We propose that the role of vOT in familiar word recognition is strictly sequential, feedforward. In contrast, feedback interaction is selectively engaged in reading unfamiliar letter strings, where lexical uncertainty probably modulates the direction of information flow for additional top-down linguistic constraints to facilitate comprehension. This adaptability likely reflects the dynamic interplay of hierarchical processing streams that constrain and optimize word reading.

## 5 Data availability statement

The MEG and MRI data are available upon reasonable request from the authors; the data is not publicly available as it contains personal information, and its reuse for other research purposes requires a new ethical review. Numerical data for the figures, which have been highly processed to prevent identification of individual participants, is available at https://osf.io/yzqtw.

## 6 Code availability statement

For connectivity analysis, we used MNE-Connectivity package (https://github.com/mne-tools/mne-connectivity). The custom code used in the study can be accessed at https://github.com/AaltoImagingLanguage/you2025.

## 7 Acknowledgments

This work was funded by the Academy of Finland (#346585 and #343385 to M.v.V., #355407 to R.S.), the Sigrid Jusélius Foundation (to R.S.), the Medical Research Council (UK: MC UU 00030/9 to O.H.), and the China Scholarship Council (#202206100022 to J.Y.). For the purpose of open access, the UKRI-funded author O.H. has applied a Creative Commons Attribution (CC BY) license to any Author Accepted Manuscript version arising from this submission. The authors thank Aino Saranpää, Mia Illman, and Rami Kunnas for their assistance with data collection. The authors acknowledge the computational resources provided by the Aalto Science-IT project.

## 8 Author contributions

Conceptualization: J.Y., O.H., R.S., M.v.V.; Data collection: J.Y.; Methodology: J.Y., O.H., M.v.V; Interpretation: J.Y., O.H., R.S., M.v.V.; Writing—original draft: J.Y.; Writing—review & editing: J.Y., O.H., R.S., M.v.V.

