## Supplemental Figures for "Separating feedforward and feedback dynamics using time-frequency-resolved connectivity: a hybrid model of left ventral occipitotemporal cortex in word reading"

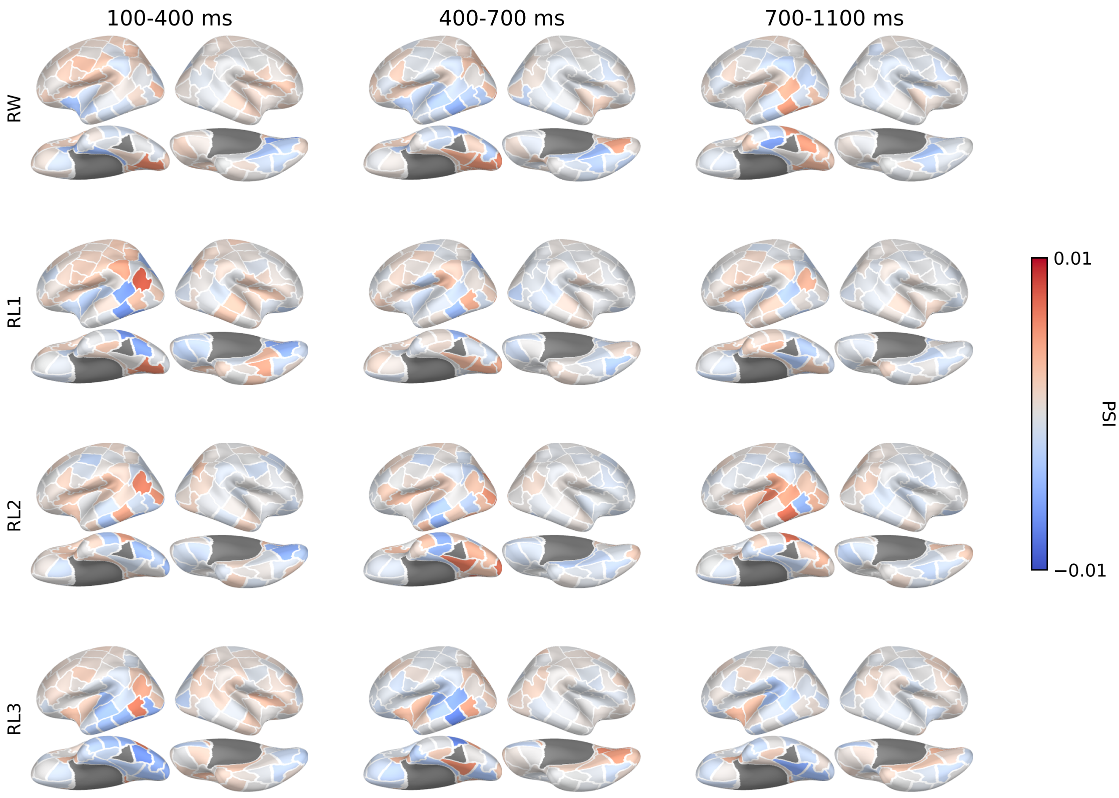


Figure S1: Group-level whole-cortex directed connectivity patterns in the three experimental conditions and three time windows, with the left vOT as seed region and analyzed using PSI in the theta band (4–7 Hz).

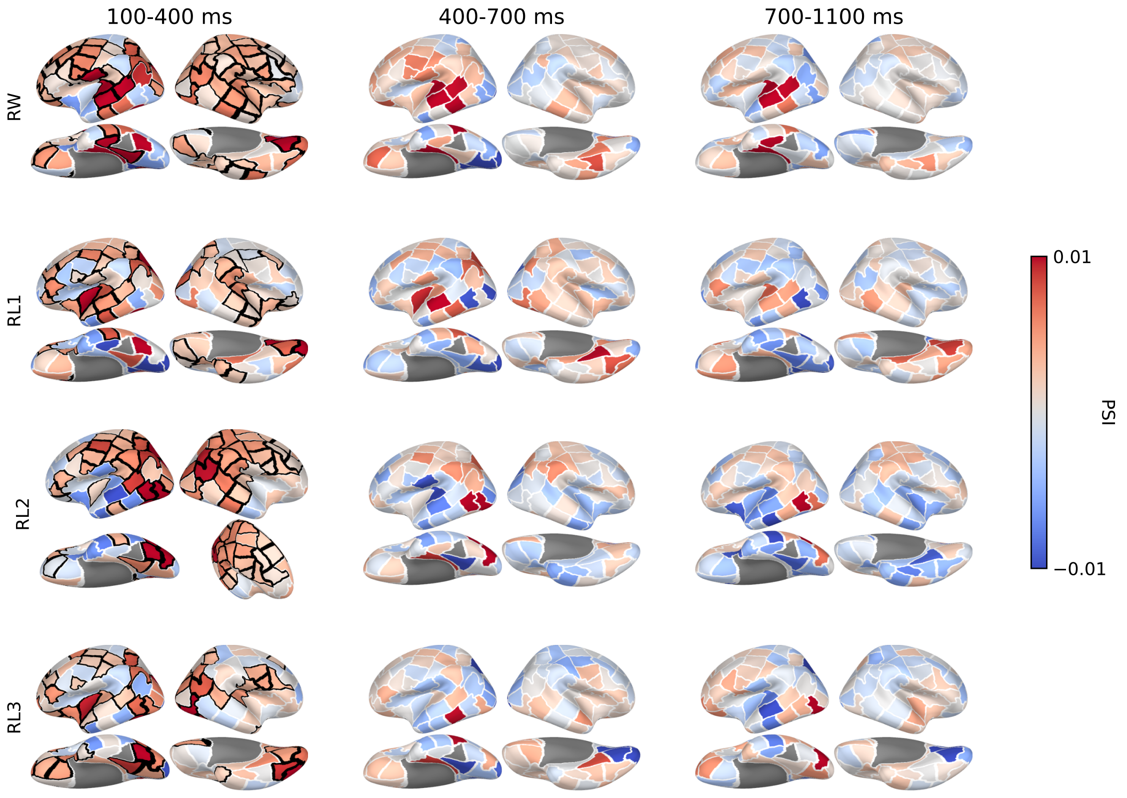


Figure S2: Group-level whole-cortex directed connectivity patterns in the three experimental conditions and three time windows, with the left vOT as seed region and analyzed using PSI in the alpha band (7–13 Hz).

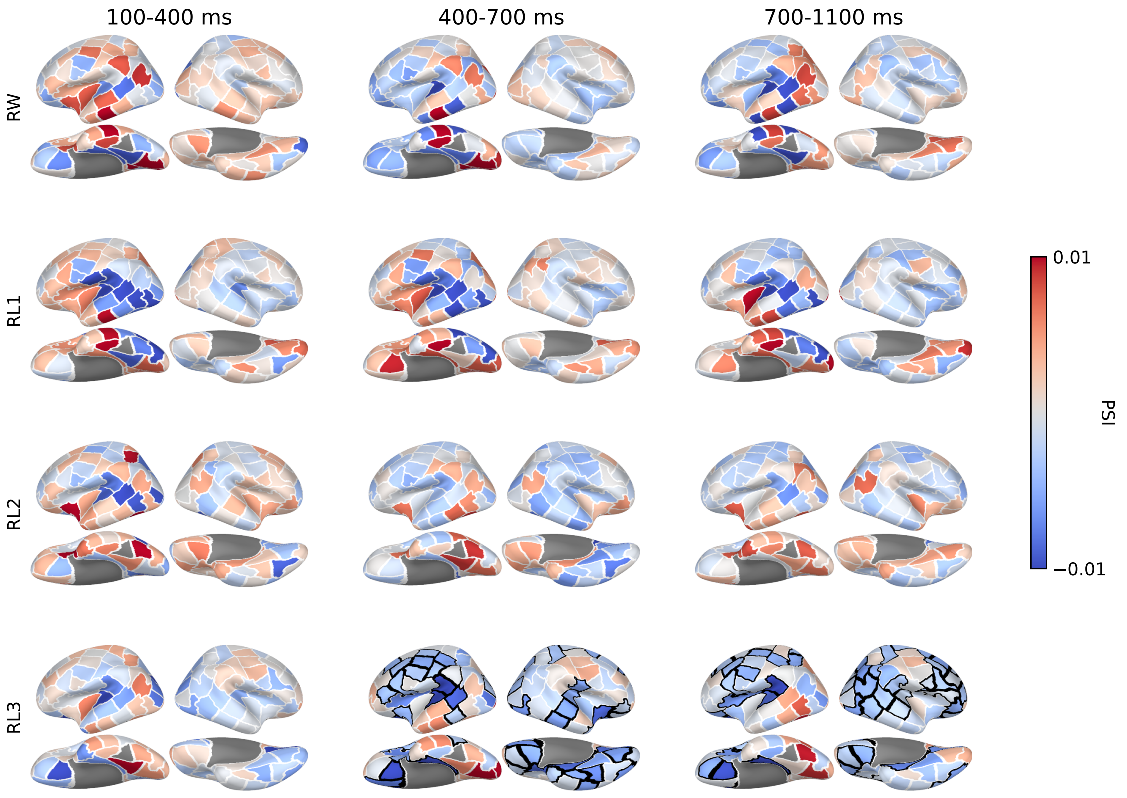


Figure S3: Group-level whole-cortex directed connectivity patterns in the three experimental conditions and three time windows, with the left vOT as seed region and analyzed using PSI in the low-beta band (13–20 Hz).

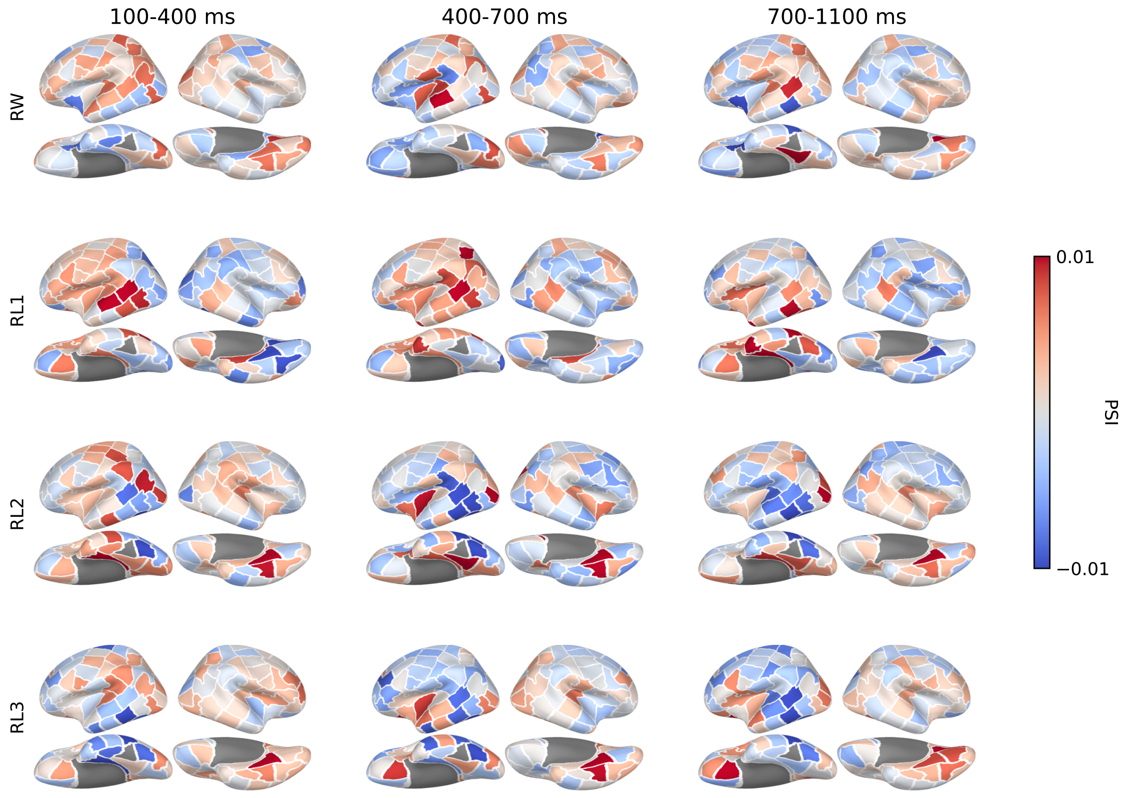


Figure S4: Group-level whole-cortex directed connectivity patterns in the three experimental conditions and three time windows, with the left vOT as seed region and analyzed using PSI in the high-beta band (20–30 Hz).

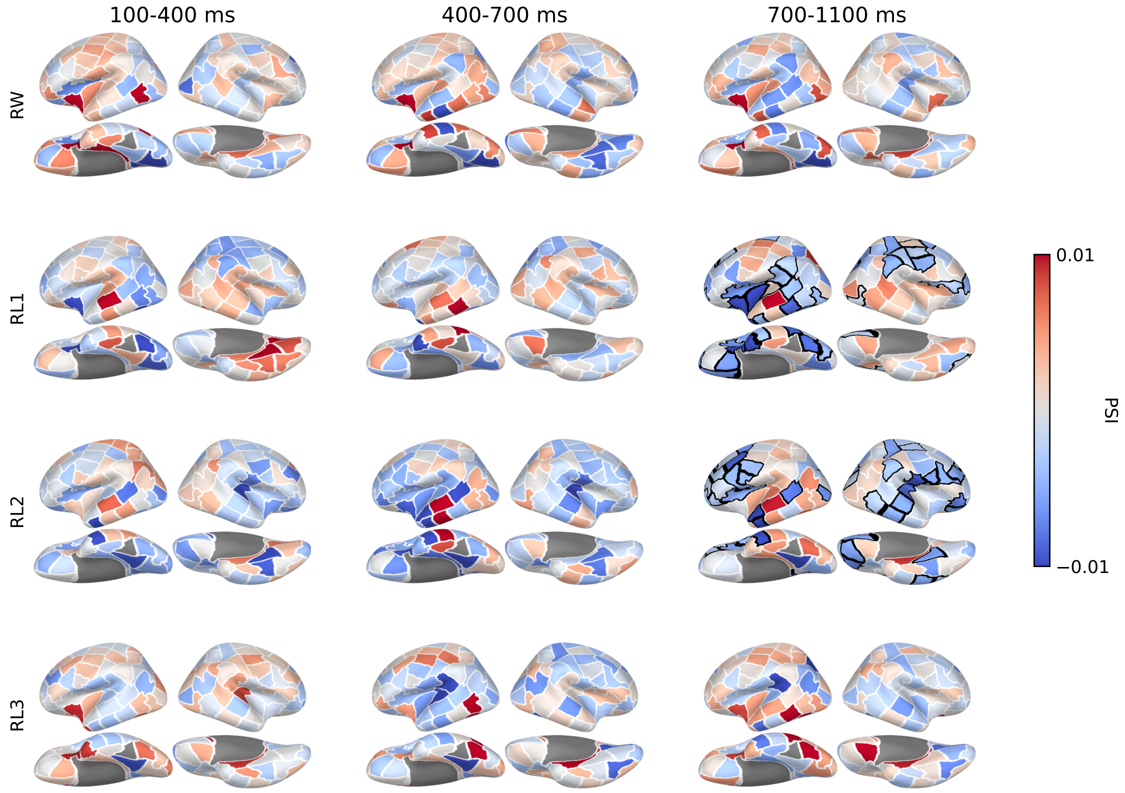


Figure S5: Group-level whole-cortex directed connectivity patterns in the three experimental conditions and three time windows, with the left vOT as seed region and analyzed using PSI in the low-gamma band (30–40 Hz).

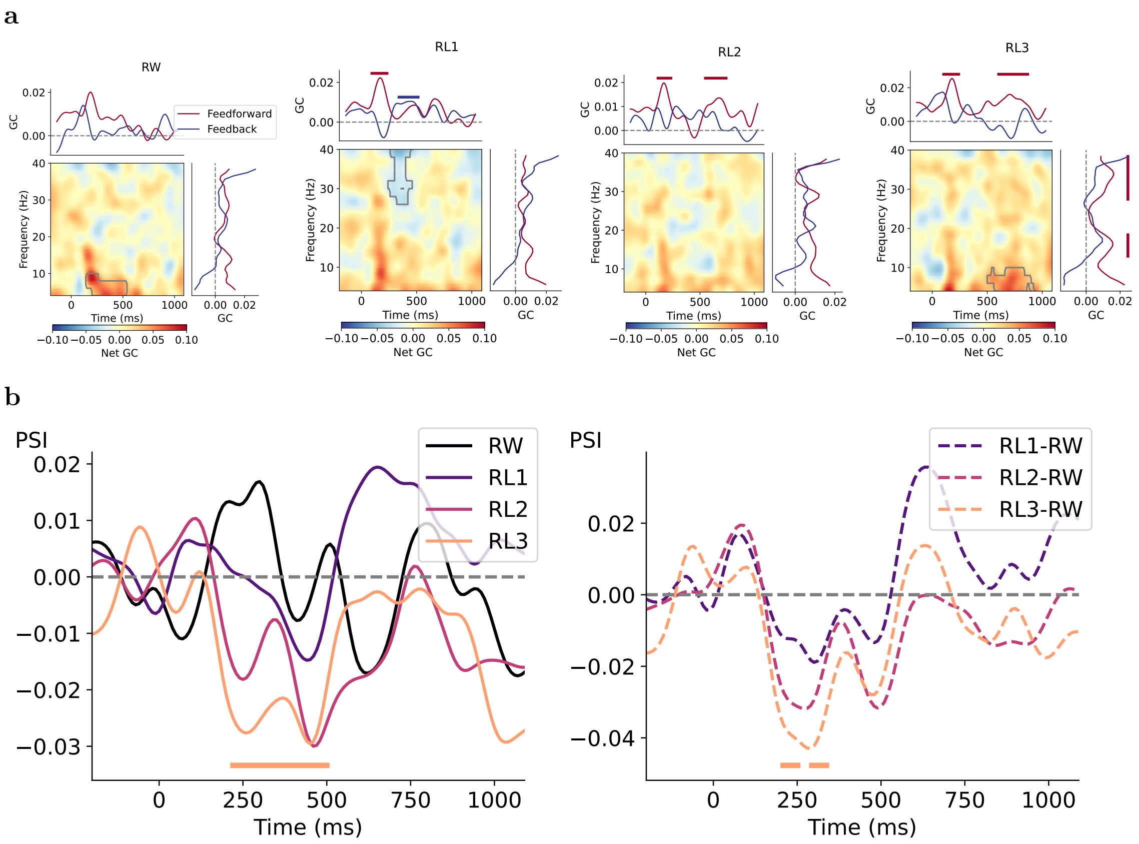


Figure S6: Time-varying causal influences between left AT and vOT for each condition. **a**, In the subplot for each condition, left-bottom panel displays the time-frequency net GC map, in which significant clusters with positive values (red) indicate dominant feedforward connectivity (vOT $\rightarrow$ AT) and negative values (blue) indicate dominant feedback connectivity (AT $\rightarrow$ vOT). Curves on top and right represent separate feedforward and feedback Granger causal influence averaged over frequency and time, respectively. Gray borders within the maps and solid bars above the plots indicate time-frequency/time/frequency clusters associated with $p < 0.05$ based on cluster-based permutation tests against zero. **b**, Temporal dynamic of PSI values for each condition (left) and their contrasts between condition pairs (right). Colored bar under the plot denotes the time cluster associated with $p < 0.05$ based on two-tailed cluster-based permutation tests.

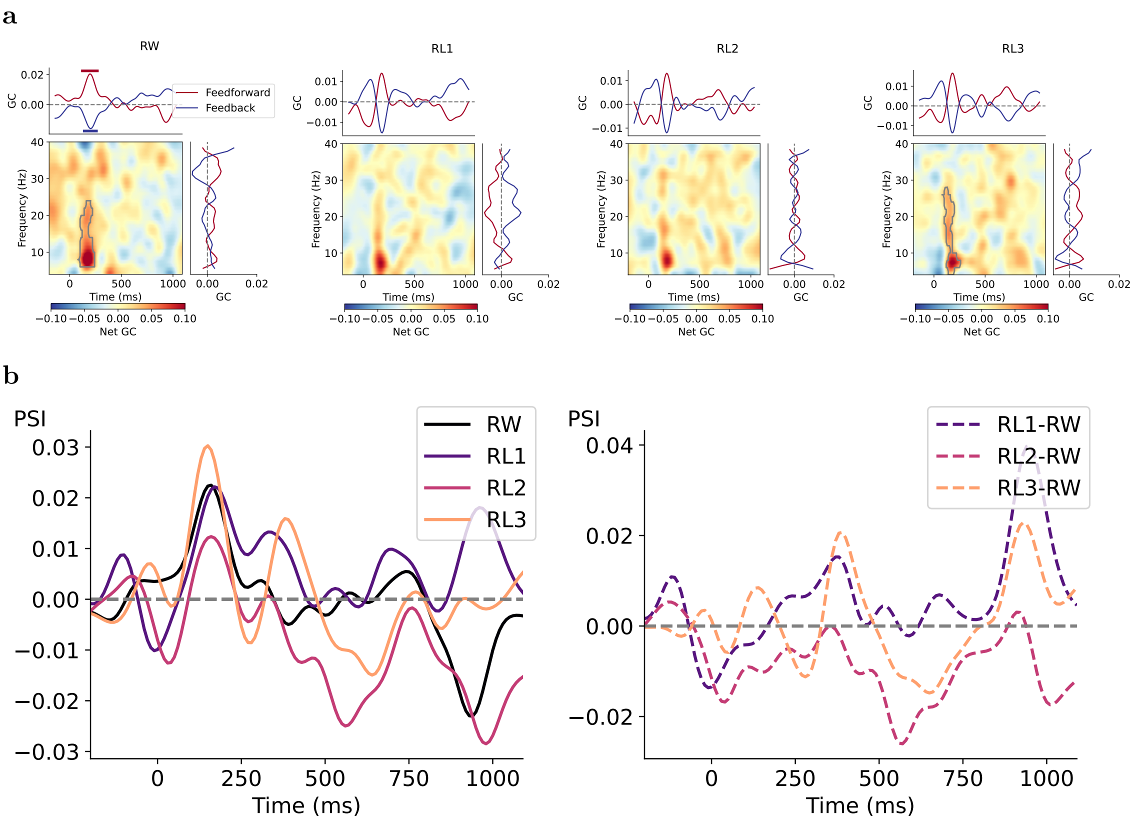


Figure S7: Time-varying causal influences between left pC and vOT for each condition. **a**, In the subplot for each condition, left-bottom panel displays the time-frequency net GC map, in which significant clusters with positive values (red) indicate dominant feedforward connectivity (vOT $\rightarrow$ pC) and negative values (blue) indicate dominant feedback connectivity (pC $\rightarrow$ vOT). Curves on top and right represent separate feedforward and feedback Granger causal influence averaged over frequency and time, respectively. Gray borders within the maps and solid bars above the plots indicate time-frequency/time/frequency clusters associated with $p < 0.05$ based on cluster-based permutation tests against zero. **b**, Temporal dynamic of PSI values for each condition (left) and their contrasts between condition pairs (right). No time clusters are associated with $p < 0.05$ based on two-tailed cluster-based permutation tests.
